# Evolutionary conditioning enables guided generation of functionally diverse enhancers

**DOI:** 10.64898/2026.04.13.718170

**Authors:** Andrew G. Duncan, Micaela E. Consens, Lorin Crawford, Jennifer A. Mitchell, Alan M. Moses, Kevin K. Yang, Alex X. Lu

## Abstract

Deep learning has been instrumental in our understanding of how enhancers encode regulatory information in their DNA sequence and has demonstrated preliminary success with enhancer design. However, the prevailing approach for enhancer design, cell type label conditioning, depends on labeled data from massively parallel reporter assays, which only exists for a handful of cell types. We propose EnhancAR, an autoregressive model trained on sets of unaligned homologous enhancer sequences to learn the function of the enhancer conserved over evolution and generate sequences that resemble real homologs. By training EnhancAR on 1.7 million human enhancer homolog sets spanning 1,888 cell types, EnhancAR generates enhancers for a variety of contexts without being conditioned on a cell type label. We computationally validate that when conditioned on a set of enhancer homologs, EnhancAR generates novel and diverse sequences that preserve the functional properties of the homologs. By prompting EnhancAR with homologs for existing cell type specific enhancers, we design enhancers with similar predicted cell type specificity. We further demonstrate that when trained on length sorted homologs, EnhancAR can design enhancers shorter than the conditioning homologs that preserve the predicted activity. In summary, we find that leveraging evolutionary information in enhancer homologs enables a more flexible and general paradigm for designing enhancers with specific functions.

## Introduction

Enhancers are regulatory elements that control the spatiotemporal expression of genes and are essential for development and evolution (Jindal & Farley, 2021; Long et al., 2016; Spitz & Furlong, 2012). The ability to design enhancers with specific functions, such as enhancers that drive gene expression in a specific cell type, is important for applications in gene therapy (Artemyev et al., 2024) and synthetic biology (Brewster & Parisutham, 2026). However, the rational design of enhancers is difficult, as the cis-regulatory code, which specifies how enhancer function is encoded into DNA sequence, remains poorly understood and is a major unanswered question in biology (Smith et al., 2023; Zeitlinger et al., 2024). Although enhancers are broadly known to contain multiple binding sites that enable interactions with transcription factors, how the composition and positioning of binding sites ultimately lead to function remains unclear, in part due to complex synergistic interactions between recruited transcription factors.

Motivated by the need to model the cis-regulatory code without prior knowledge, researchers have trained deep learning models on data assaying the activity of enhancer sequences (de Almeida et al., 2022; Kelley et al., 2016; Zhou & Troyanskaya, 2015). These studies typically use plasmid-based massively parallel reporter assays (MPRA) that measure enhancer activity for tens of thousands of sequences within specific cell types, meeting the demand for data required to train deep learning models. Often, these are sequence-to-function models, trained to predict the assay measurements from DNA sequence. Once trained, post-hoc analyses (Koo et al., 2021; Shrikumar et al., 2020) have demonstrated that these models learn aspects of the cis-regulatory code (de Almeida et al., 2022; Kelley et al., 2016), which can be used for enhancer design through approaches like sequence evolution and motif implanting (Castillo-Hair et al., 2025; Chen et al., 2025; de Almeida et al., 2022; Gosai et al., 2024; Taskiran et al., 2024). More recent work aims to train generative deep learning models to design enhancers (DaSilva et al., 2026; Lal et al., 2024; Sarkar et al., 2024; Taskiran et al., 2024), which directly generate DNA sequence.

As generating enhancers that are functional in specific contexts is a common motivation of generative models, and because enhancer data is still predominantly from cell type specific MPRAs, current generative models implement cell type label conditioning (DaSilva et al., 2026; Sarkar et al., 2024): given a cell type label, the model generates enhancer sequences that are likely to have activity within that cell type. However, training these models requires data from experimental assays where activity is measured in a cell type, limiting training and inference to conditions where data is available and biasing for conditions that are experimentally feasible (e.g., cell lines amenable to experimental culture). Moreover, it is unclear whether models trained on assay data from one context will generate enhancers that are specific to that context or may show expression more broadly, or how well cell-based assays reflect expression *in vivo*. Here we seek to explore generative enhancer models that do not have the inherent limitations of explicit cell type conditioning.

Our approach to generative modeling of biological sequences uses homologs (sequences related through evolutionary descent) as context to prompt language models to generate similar sequences (Truong & Bepler, 2023; Xie et al., 2025; Yang et al., 2025). This approach leverages the key idea that evolution conserves the function of non-coding DNA while allowing the sequence encoding that function to diverge. Thus, rather than conditioning models through a single lab assay, or a set of assays, to define function, conditioning on homologs potentially allows for the generation of sequences with a specific function that has been preserved through evolution. As this approach has been successfully applied to other biological sequences such as proteins (Truong & Bepler, 2023; Yang et al., 2025) and promoters (Xie et al., 2025), we reasoned that it would be applicable to enhancers as well. We trained EnhancAR, an autoregressive generative model, on sets of unaligned homologous enhancer sequences. At inference time, this allows us to perform conditional design of enhancers by prompting EnhancAR with the homologous sequences of a specific enhancer. Using a combination of *in silico* evaluations, we confirm that EnhancAR uses prompts to guide the generation of sequences with similar functions. We further demonstrate the utility of EnhancAR for downstream applications, including cell type specific enhancer design that does not require cell type labels and shortening enhancers while preserving function.

## Results

### EnhancAR is an autoregressive model for enhancer generation

To train a model that generates enhancer sequences based on prompted homologous enhancer sequences, we assembled, to our knowledge, the largest dataset of homologous enhancer sequences to date (Figure 1A). We first downloaded candidate human enhancers from ENCODE SCREEN (ENCODE Project Consortium et al., 2020; Moore et al., 2026), which contains over 1.7 million candidate enhancers across 1,888 human cell types defined using a standard set of functional measurements. We used the coordinates of these enhancers in the human genome to extract homologous sequences from other mammals in the Zoonomia 241 mammalian species whole-genome alignment (Zoonomia Consortium, 2020). In total, we identified 233,158,475 homologs, for an average of 135 homologs per enhancer. We define an enhancer homolog family as an unaligned set of enhancer sequences with shared ancestry that are identified through whole-genome alignments. We held out enhancer homolog families where the human sequence was on chromosomes 3 and 19 for the test set and 1,000 random enhancer families from the training set for validation. The validation set was only used for assessing model convergence and losses, whereas the test set was used for post-training evaluation.

**Figure 1.**
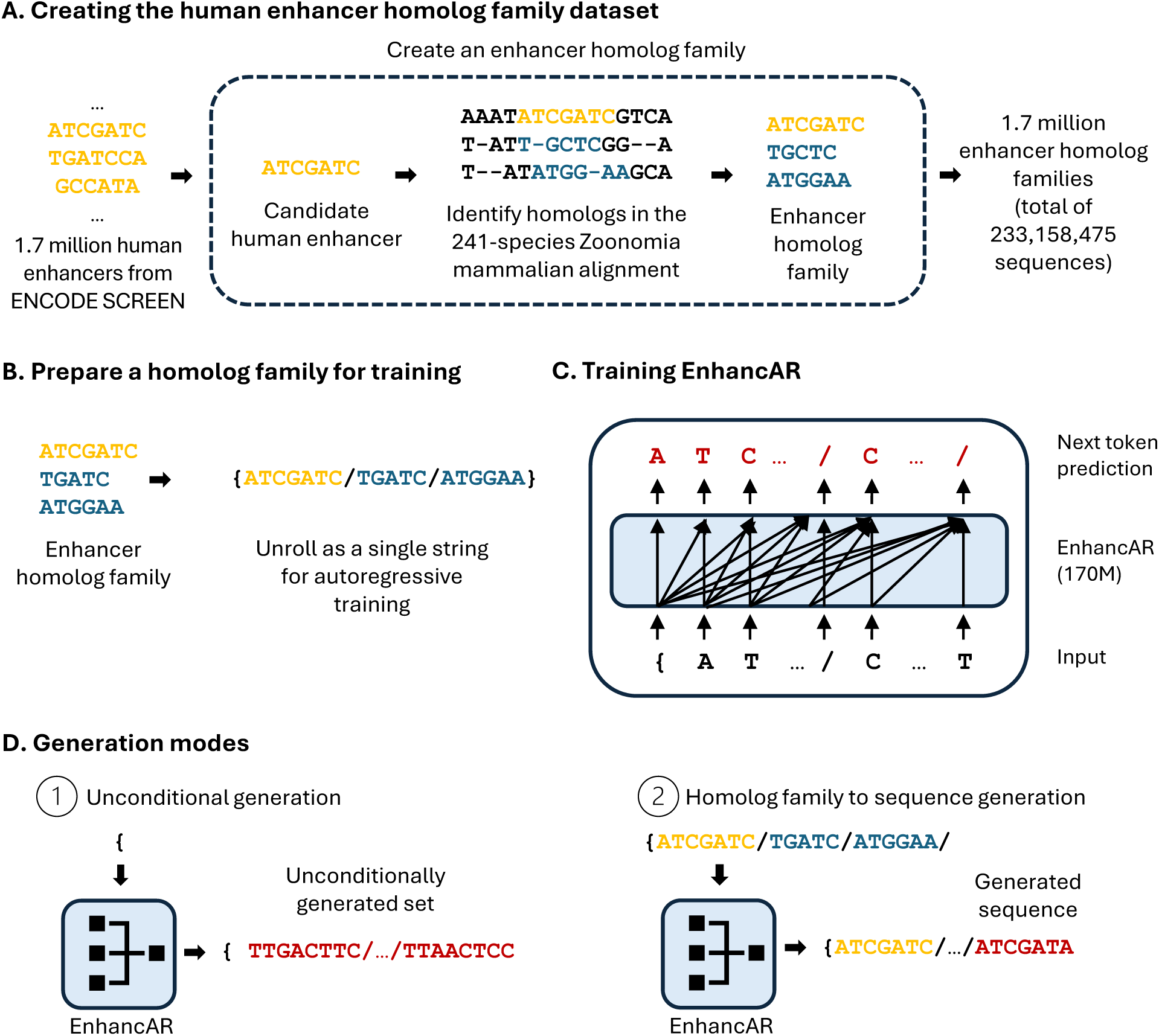
Data preparation, model training, and sequence generation with EnhancAR. (A) EnhancAR was trained on over 1.7 million enhancer homolog families for candidate human enhancer sequences from ENCODE SCREEN. (B) Enhancer homolog families are prepared for training by unrolling into a single string of characters separated by special characters. (C) EnhancAR (170M parameters) is trained with next token prediction on tokenized homolog families. (D) At inference time, EnhancAR has a variety of generation modes. EnhancAR can generate a sequence set unconditionally (1) or extend a homolog family (2).

We used this dataset to train EnhancAR, an autoregressive model of enhancer evolution. Since EnhancAR is trained on enhancer homolog families, we reasoned this would enable the model to generate novel sequences that share function with real conditioning sequences. We chose a Jamba model (Lieber et al., 2024), which is a hybrid transformer/state-space model, allowing the model to use longer context windows with lower memory usage than transformer architectures (Vaswani et al., 2023), which we reasoned would enable EnhancAR to learn features across long chains of homologs.

To prepare each enhancer homolog family for training, we sampled 512 base pair (bp) regions from up to 64 homologous enhancer sequences and constructed a sequence of sequences separated by special characters (see Methods) (Figure 1B). We trained two versions of EnhancAR: one trained on randomly ordered homologs (EnhancAR) and one trained on homologs sorted by descending length (EnhancAR-sorted). EnhancAR was trained with a standard next-token prediction task with cross-entropy loss, using the previous ground truth tokens as context (Figure 1C). This training procedure results in a model that generates a probability distribution predicting the next token in a sequence. During inference time, we can generate new sequences by iteratively generating one token at a time, until a complete sequence or set of sequences is generated. We experiment with two modes of generation (Figure 1D). First, as a basic validation of the model’s ability to learn the natural distribution of enhancers, we generate sequences from scratch (i.e., unconditional generation, with no homolog prompts). Second, we generate single sequences using enhancer homolog families as the starting point (homolog family to sequence generation).

### Unconditional generations from EnhancAR resemble enhancer homolog families

We first investigated EnhancAR’s ability to generate sets of sequences unconditionally (without any sequence prompts) to determine whether unconditionally generated sets resemble real enhancer homolog families. We chose to investigate unconditionally generated sets rather than individual unconditional generations, as individual enhancer sequences have a low signal-to-noise ratio in proxy metrics for function like matches to transcription factor binding site motifs, making interpretation challenging. Since our model natively generates sets of enhancers with putative homology, we took advantage of this to test if EnhancAR is capturing signals that are robustly conserved over evolution (and thus likely to be functional). If so, we reasoned that unconditionally generated sequence sets would share similarity in both sequence, motif, and functional space.

We used EnhancAR to unconditionally generate 1,000 sequence sets composed of 63 sequences. We first confirmed that the model was not copying sequences from the training dataset by conducting a nucleotide BLASTN (Sayers et al., 2025) search of the generated sets against the training dataset. As a comparison, we also conducted a BLASTN search on 1,000 held-out homolog families from the test set against the training dataset. We observed that less than 1% of the unconditional generated sequences and held-out homolog families had a significant match to the training sequences (see Methods). These results demonstrate that EnhancAR did not reproduce the training dataset for unconditional generation.

Next, we investigated the diversity of unconditionally generated sequences by measuring how similar sequences within each sequence set were to each other. Due to shared ancestry, enhancer homologs are expected to have greater overall sequence similarity than in unrelated sequences. We measured the diversity of each sequence set by calculating the average pairwise percent identity across all sequences within each set (Figure 2A). To define the expected level of conservation for enhancers across mammals, we also analyzed the 1,000 homolog families from the held-out test set. To define a lower bound of conservation, we analyzed the 1,000 held-out homolog families after they were scrambled, where nucleotides within each sequence are randomly shuffled, likely destroying the function. The generated sequence sets had lower mean sequence similarity than the held-out homolog families (Mann-Whitney U test, U=206502, p-value=2.34e-114), with a mean sequence similarity of 65.32% (Standard Deviation (SD) 3.37) for the unconditional generations (Figure 2B, blue bar) and 73.66% (SD 1.50) for the held-out homolog families (Figure 2B, orange bar). Both generated sets (Mann-Whitney U test, U=851740, p-value=6.76e-163) and held-out homolog families (Mann-Whitney U test, U=999997, p-value=0) had higher mean sequence similarity than the scrambled sequences (mean 57.05%, SD 0.92) (Figure 2B, green bar). These results suggest that EnhancAR captures some of the relationship of sequence similarity among sequences when generating sequence sets unconditionally, though there is lower sequence similarity than within the held-out homolog families.

**Figure 2.**
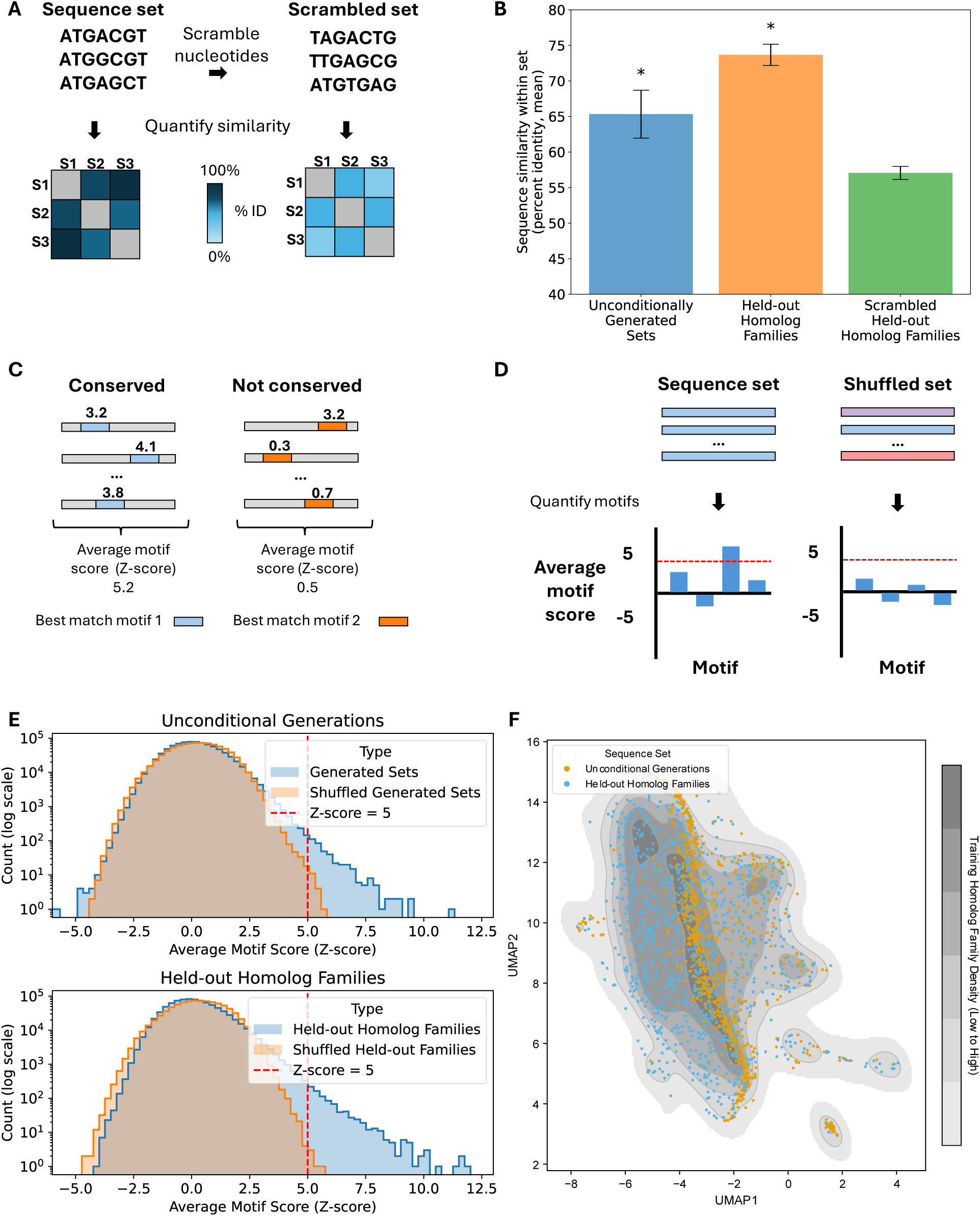
Unconditional generated sequence sets resemble natural enhancer homolog families. (A) Schematic for calculating the diversity of a sequence set and a scrambled nucleotide sequence set. (B) Bar plot showing the sequence similarity within sets, measured as the mean percent identity, of the EnhancAR unconditionally generated sequence sets (n=1,000), held-out homolog families from the test set (n=1,000), and scrambled (nucleotide shuffled) held-out homolog families (n=1,000). Error bars represent the standard deviation of the mean percent identity. Significance was assessed by comparing groups to the scrambled homolog families using the Mann-Whitney U test. Asterisks represent significance (p-value < 0.05) after Benjamini-Hochberg correction. (C) The average motif score calculation for a conserved and not conserved motif in a homolog family. Grey rectangles are homolog sequences, blue rectangles are best matches to motif 1, and orange rectangles are best matches to motif 2. (D) Schematic for quantifying motifs in sequence sets compared to shuffled sets. Light blue rectangles are homolog sequences from set A, light purple rectangles and light red rectangles are homolog sequences from randomly selected homolog families. (E) Histograms showing the normalized average motif scores for the unconditional generation sets (top) and held-out homolog families (bottom). Generated and held-out test sets are shown in blue (n=1,000), and shuffled sets are shown in orange (n=1,000). A red vertical line is placed at an average motif score of 5. Bin widths are set to 0.25. (F) Scatter plot of the Enformer embeddings UMAP of training data (grey kernel density plot) (n=50,000), unconditionally generated sequences (yellow points) (n=1,000), and held-out homolog family sequences (blue points) (n=1,000). The UMAP was fit to training dataset, and the unconditional generations and held-out homolog family sequences were transformed using the UMAP model.

We then examined whether the unconditionally generated sequence sets resembled held-out homolog families in motif space. As enhancer homologs conserve binding sites for functionally important transcription factors (Wong et al., 2020), we hypothesized that the unconditionally generated sets would similarly maintain binding sites, which we could measure by analyzing if known motifs corresponding to binding sites are maintained. We first measured motif composition by scanning each sequence in each set with the 879 motifs from the JASPAR 2024 non-redundant vertebrate motif set (Rauluseviciute et al., 2024). For each generated set, we then computed an average motif score (AMS) for each motif by averaging the highest score across each sequence (Figure 2C), as averaging motif scores over homologs have been shown to improve the signal-to-noise ratio for motif matches (Alam et al., 2023). As motifs vary in length and composition, the range of match scores can vary. To put the scores on the same scale, we normalized the AMSs by repeating the calculations on 100 scrambled versions of each motif and converted the raw AMS to a Z-score (see Methods) (Alam et al., 2023). We used a Z-score cutoff of 5 to identify matches that are more than five standard deviations above the expectation obtained from scrambling motifs, as this was near the maximum AMS seen in shuffled held-out homolog families.

We compared the AMSs for the 1,000 generated sets and the 1,000 held-out homolog families, counting motifs in score bins (Figure 2D). As a baseline, we also quantified motif enrichment for shuffled sets, where sets are formed from distinct enhancers sampled at random from the test dataset (as opposed to homologs). As these enhancers likely span a variety of functions, we reasoned that this estimates how much motif enrichment is expected by random chance. We observed that both generated and held-out homolog families contained many enriched motifs with AMS greater than 5 (Figure 2E, blue histogram), whereas shuffled sets contained very few (Figure 2E, orange histogram). Taken together, these results demonstrate that sequence sets generated unconditionally by EnhancAR share sequence and motif similarity that resemble held-out enhancer homolog families.

Having demonstrated that the generated sets resembled held-out homolog families, we next investigated whether these generated sequences covered the functional space of human enhancers. To measure the function of a sequence, we used embeddings generated by Enformer (Avsec et al., 2021), a deep learning model trained to predict thousands of genomic tracks from DNA sequence (see Methods). We projected the Enformer embeddings for unconditionally generated sequences and held-out homolog sequences onto a UMAP of Enformer embeddings for 50k random training set homolog sequences (Figure 2F). We find that the held-out homolog families (Figure 2F, blue dots) broadly cover the functional space of the training data (Figure 2F, grey density). The unconditionally generated sequences (Figure 2F, yellow dots) also cover a range of the functional space; however, they tend to stay near the mode. As the motif matches present in an enhancer encode the function of an enhancer, we repeated this experiment on a UMAP of AMSs calculated across sequence sets, observing a similar pattern (Supplementary Figure 1).

### EnhancAR generates novel and diverse sequences that preserve the motif composition of their prompts

Having established that our model could unconditionally generate sequence sets that resembled our held-out enhancer homolog families, we next wanted to understand if EnhancAR, when prompted with sequences from an enhancer homolog family, was capable of generating sequences with similar functional and composition properties. To explore this, we used the previously sampled 1,000 held-out enhancer homolog families from our test set, sampled up to 63 sequences from each set (see Methods), and generated 63 new sequences with the same prompt. As we observed that EnhancAR had a tendency to not terminate generation at the expected length (Supplementary Figure 2), we post-processed generations by trimming from the right side until they reached the length of the longest sequence in the prompt.

As we expected that the generated sequences would resemble their prompts in sequence space, we measured how similar generated sequences were to their prompts, analyzing one generation per prompt. We calculated the pairwise percent identity between each generated sequence and their prompt sequences and obtained the value for the most similar sequence as a measure of novelty (Figure 3A). For comparison, we measured the sequence similarity of three sequence types: human sequences, scrambled human sequences, and prompt-derived consensus sequences. Human sequences were held out during generation, providing an expectation for how similar a true homolog might be to the prompt. To establish a baseline representing sequences with no functional relationship to the prompt, we scrambled the human sequences. Our second baseline was the prompt-derived consensus sequence, which represents a naïve design strategy. We found that the generated sequences had lower sequence similarity with the prompts than the human sequences had with the prompts, with mean sequence similarity of 86.60% (SD 7.79) in generated sequences (Figure 3B, blue bar) compared to 98.26% (SD 1.70) in the human enhancer sequences (Figure 3B, orange bar). In contrast, the scrambled sequences (Figure 3B, red bar) had significantly lower sequence similarity (Mann-Whitney U test, U=991653.5, p-value=0) than the generated sequences (Figure 3B, blue bar), with a mean sequence similarity of 60.05% (SD 3.23).

**Figure 3.**
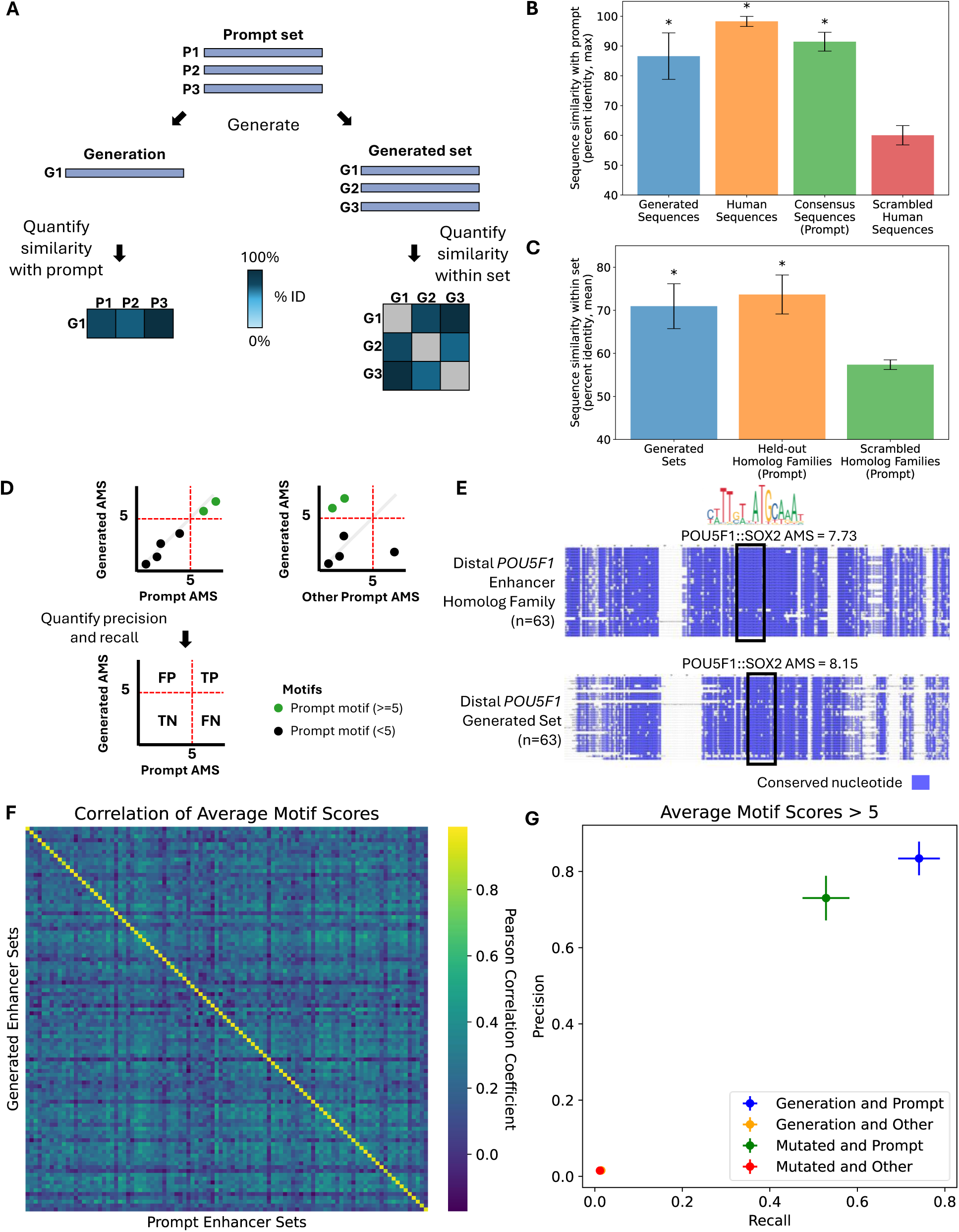
Test set generations resemble their prompts in sequence and motif space. (A) Novelty measures how similar a generated sequence is to the prompt. Diversity measures how similar generated sequences from the same prompt are (sequence set). (B) Bar plot showing the sequence similarity between sequences and their prompt, measured as the maximum percent identity, of EnhancAR generated sequences (n=1,000), human sequences (n=1,000), and scrambled human sequences (n=1,000) for held-out homolog family prompts (n=1,000). Significance was assessed by comparing groups to the scrambled human control using the Mann-Whitney U test. Asterisks represent significance (p-value < 0.05) after Benjamini-Hochberg correction. Error bars represent the standard deviation of the maximum percent identity. (C) Bar plot showing the sequence similarity between sequences within a set, measured as the mean percent identity of the EnhancAR generated sequences (n=1,000), the prompt sequences (n=1,000), and scrambled prompt sequences (n=1,000) for held-out homolog family prompts (n=1,000). Significance was assessed by comparing groups to the scrambled homolog family using the Mann-Whitney U test. Asterisks represent significance (p-value < 0.05) after Benjamini-Hochberg correction. Error bars represent the standard deviation of the mean percent identity. (D) Average motif scores are computed for prompt sequence sets and generated sequence sets. The correlation of average motif scores between the different sets was then calculated. Finally, the precision and recall for motifs with an average motif score > 5 were calculated between sets. (E) Multiple sequence alignment (MSA) for the 63 prompt sequences from the *POU5F1* enhancer homolog family (top) and 63 generated sequences using the *POU5F1* enhancer family prompt sequences (bottom). The black box denotes the region of the alignment corresponding to a strongest POU5F1::SOX2 JASPAR motif (MA0142.1) match. The blue represents conserved nucleotides in the alignment. The average motif scores for the POU5F1::SOX2 motif is shown for each sequence set. (F) Heatmap of the average motif score similarity (Pearson correlation) between each generated sequence set (n=100) and each prompt set (n=100) for a randomly sampled subset of the held-out homolog families (n=100). (G) Scatter plot of the precision compared to recall of average motif scores > 5 for the generated sequence sets with their own prompt and all other prompts using the held-out homolog families with at least 2 motifs with an average motif score > 5 (n=176). As a control, the precision and recall for mutated prompt sets (mutation rate 15%) compared to their own prompt and all other prompts. Vertical lines represent two times the standard error of precision and horizontal lines represent two times the standard error of recall.

Next, we evaluated the sequence diversity of the generated sequence sets. We reasoned that if the generated sequences resembled true homologs of their prompts, then if we grouped multiple generations from the same prompt into a set, the set would have similar average sequence similarity as their prompt. For comparison, we also measured the diversity of scrambled sequence prompts, which represent a baseline for the sequence similarity of unrelated sequences. We observed a mean sequence similarity of 70.95% (SD 5.22) in the generated sequences (Figure 3C, blue bar), which was similar to the mean sequence similarity of 73.66% (SD 4.53) in the prompt sequences (Figure 3C, orange bar). By comparison, the mean sequence similarity of 57.36% (SD 1.13) in the scrambled homolog families (Figure 3C, green bar) was significantly lower than both the generated sets (Mann-Whitney U test, U=999540.0, p-value=0) and held-out homolog families (Mann-Whitney U test, U=999997.0, p-value=0). Altogether, these results demonstrate that EnhancAR is generating novel and diverse sequences that resemble true homologs of the prompt sequences.

We then investigated whether the generated sequences preserved the motif composition of their prompt. We used the AMSs across sequence sets to represent the enrichment of transcription factor motifs in each sequence set. We looked at the overall motif similarity, using the Pearson correlation to measure the similarity of the AMS representation between each generated sequence set with all prompts (Figure 3D, top). We found that the average motif score representations for generated sequence sets had significantly higher correlation (Mann-Whitney U test, U=990000, p-value=7.5e-67) with their prompts (average correlation 0.9376, standard error (SE) 0.0031) than to other prompts (average correlation 0.2167, SE 0.0012) (Figure 3F), demonstrating that the generations are specific to their prompt.

For each enhancer homolog family, only a handful of the 879 JASPAR vertebrate transcription factor motifs are enriched and relevant for enhancer function. To focus on these strongly enriched motifs, we selected the motifs in each enhancer homolog family with an AMS greater than 5, the same cut-off we had used for the unconditional generation analysis. As an illustrative example, we prompted EnhancAR with 63 homologs for the *POU5F1* enhancer (Figure 3E, top). *POU5F1* encodes a key pluripotency transcription factor that self-regulates by co-binding with SOX2 at the distal *POU5F1* enhancer (Chew et al., 2005). Of the 879 JASPAR motifs, only the POU5F1::SOX2 co-motif had an AMS greater than 5 in the homolog family, which was maintained in the generated set (Figure 3E, bottom). Next, as a more systematic evaluation beyond our single illustrative example, we investigated whether the test set prompted generations were preserving the strongest motifs of their prompts using precision and recall (Figure 3D, bottom). We found that on average the generated homolog families had higher precision (0.8339) (Mann-Whitney U test, U=5182679, p-value=0) and recall (0.7411) (Mann-Whitney U test, U=5173853, p-value=0) with their prompt for the strongest motifs (Figure 3G, blue point), compared to lower precision (0.0156) and recall (0.0147) with all other prompts (Figure 3G, yellow point). This is consistent with EnhancAR placing importance on the conserved motifs in the prompt and not others. In contrast, the randomly mutated prompts, with a mutation rate of 15% per sequence, displayed lower precision (0.7300) (Mann-Whitney U test, U=18308, p-value=5.8e-3) and recall (0.5284) (Mann-Whitney U test, U=20872.5, p-value=2.9e-8) with their prompt (Figure 3G, green point). Taken together, when prompted with an enhancer homolog family, EnhancAR generates sequences that maintain both overall motif similarity and the strongest motifs of their prompt.

### EnhancAR can design predicted cell type specific enhancers without cell type information at training or generation time

Having demonstrated that EnhancAR is using prompts to generate novel and diverse sequences that resemble homologs of their prompts, we next evaluated EnhancAR’s potential as a design tool for cell type specific enhancers. Unlike most generative enhancer models, EnhancAR is not explicitly conditioned on cell type. However, we reasoned we could still achieve cell type specific design without further training by prompting EnhancAR with examples from enhancer homolog families where the human enhancer has the target cell type specific activity. We selected the 500 cell type specific human enhancers for the K562, HepG2, and WTC11 cell lines (identified via MPRA (Agarwal et al., 2025)) with the largest element specificity score for each cell type (see Methods) that also had at least 64 homologs. For each human enhancer sequence, we assembled a prompt with 63 homologs (holding out the human sequence) and used the same prompt to generate 10 sequences for each set. We observed that the generations were novel compared to both the prompt and training set (Supplementary Figure 3A/C/E) and remained diverse when generated multiple times from the same prompt (Supplementary Figure 3B/D/F).

We next compared the predicted enhancer activity of the generated sequences and the prompt sequences using MPRALegNet models trained on MPRA data for each cell type (Agarwal et al., 2025) (Figure 4A). Each MPRALegNet model is a sequence-to-function convolutional neural network trained to predict enhancer activity in a specific cell-line from a 230 bp sequence. Despite no cell type information in training or prompt data, generated sequences demonstrated predicted cell type specificity and strength on par with the prompt sequences, albeit with weaker enhancer activity than the human sequences (Figure 4B). For example, the HepG2 prompted generated sequences had strong predicted activity in the HepG2 model but were no different from scrambled human sequences in the K562 and WTC11 models.

**Figure 4.**
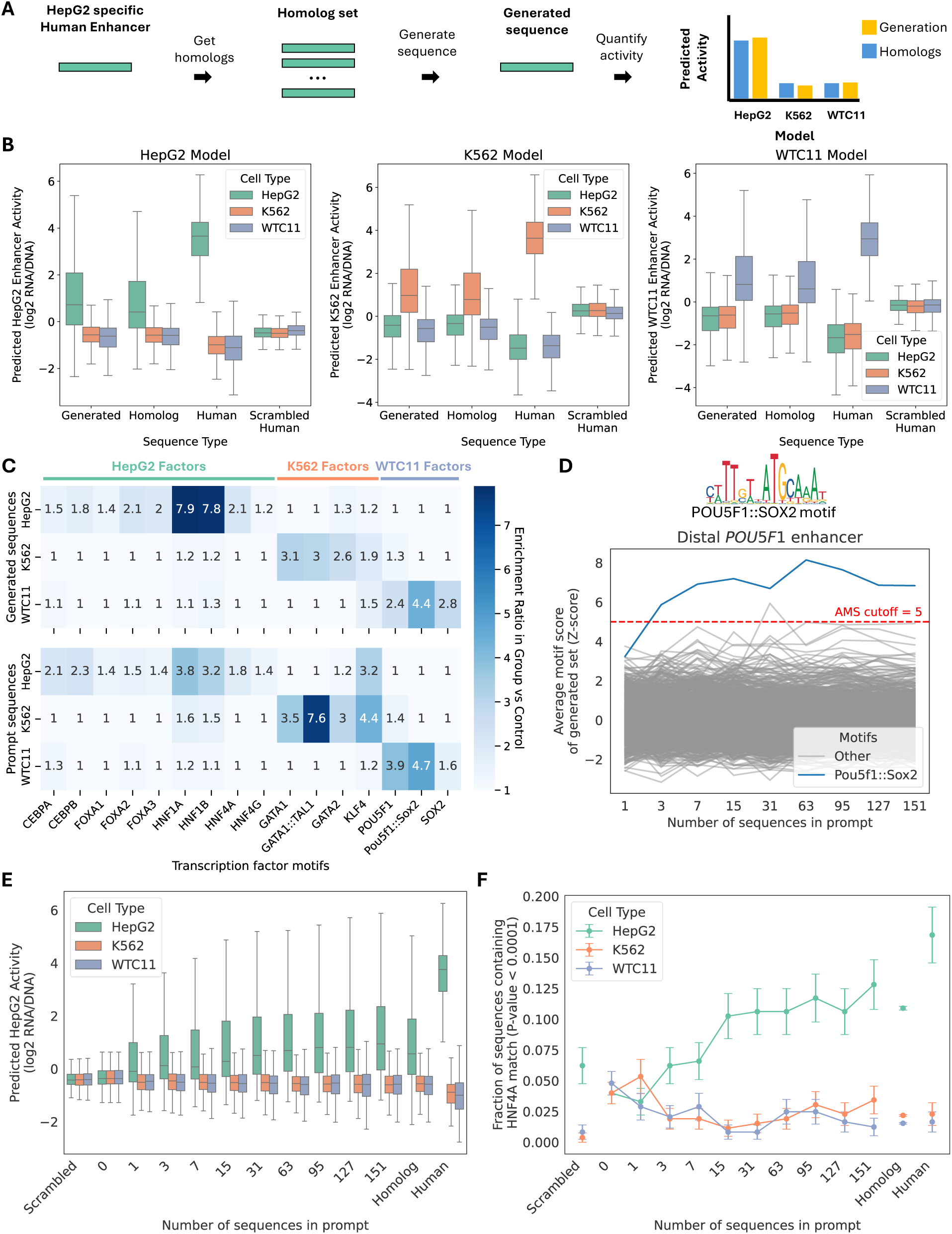
EnhancAR can design cell type specific enhancers without cell type information. (A) Cell type specific enhancers are designed by prompting EnhancAR with a homolog family for an existing cell type specific enhancer with the target cell type specificity. Cell type specificity and activity are then evaluated using the MPRALegNet models. (B) Box plot showing the predicted MPRA enhancer activity (log2(RNA/DNA)) using MPRALegNet models for HepG2 (left), K562 (middle), and WTC11 (right) for EnhancAR generated sequences, human sequences, homolog sequences, and scrambled human sequences for cell type specific HepG2 (n=500), K562 (n=500), and WTC11 (n=500) enhancer homolog families. (C) The enrichment in primary versus control sequences of cell type specific factors for HepG2 (n=500), K562 (n=500), and WTC11 (n=500) generations (top) and prompts (bottom) were calculated using the MEME-suite tool SEA. (D) EnhancAR was prompted with an increasing number of sequences in each prompt for the distal *POU5F1* enhancer homolog family. The average motif score (AMS) for 879 JASPAR 2024 non-redundant vertebrate motifs was calculated for each generated set. Motifs with AMS greater than 5 in the enhancer homolog family are highlighted. All other motifs are grey. The red dotted line represents an AMS score of 5. (E) Predicted K562 HepG2 enhancer activity (log2(RNA/DNA)) using the MPRALegNet models is shown on the y-axis for the HepG2 (n=273), K562 (n=262), and WTC11 (n=242) prompted EnhancAR designs generated with an increasing number of sequences in the prompt. (F) The proportion of HepG2, K562, and WTC11 prompted EnhancAR designs generated with an increasing number of sequences in the prompt that contain a significant match to the HepG2 specific factor HNF4A using the lightmotif package (P-value <1e-4).

We hypothesized that EnhancAR’s capacity for cell type specific generation could be explained by preserving matches to cell type specific motifs in the generations. Thus, we analyzed motif composition using known cell type specific motifs (Agarwal et al., 2025) of all sequences generated from prompts from the three cell lines using the SEA tool from MEME-suite (Bailey & Grant, 2021) and observed that generated sequences were enriched for their corresponding cell type specific motifs (Figure 4C, top). For example, HepG2 prompted generations were enriched for motifs of liver transcription factors like HNF4A but not enriched in GATA1::TAL1 or POU5F1::SOX2 motifs like those from the K562 and WTC11 cell lines. Additionally, the patterns of enrichment of cell type specific factors across generations were similar to their enrichment in the prompt sequences (Figure 4C, bottom). Overall, these results demonstrate that EnhancAR can generate novel and diverse sequences with predicted cell type specificity when prompted with cell type specific homolog sequences, and that these generations preserve similar patterns of enrichment for cell type specific factors.

We next asked how increasing the number of sequences in the prompt affected design success. We hypothesized that larger prompts would improve enhancer design by providing additional evolutionary information for EnhancAR. We generated sequences prompted with the HepG2, K562, and WTC11 enhancer homolog families with an increasing number of homologs in each prompt and predicted enhancer activity with the MPRALegNet models. We observed that as the number of sequences in the prompt increased from 0 to 63 sequences, there was an overall increase in predicted activity in the target cell type while preserving specificity (Figure 4E, Supplementary Figure 4A/C). Prompts larger than 63 sequences, which were outside the training distribution, did not improve the predicted enhancer design success (Mann-Whitney U test, P-value > 0.05). Additionally, we investigated the proportion of known cell type specific motif matches in the enhancer designs and observed an increase in the proportion of corresponding cell type specific motifs as the number of sequences in the prompt increased (Figure 4F, Supplementary Figure 4B/D).

To assess how the number of sequences in the prompt interacts with a real stem cell enhancer, we revisited the *POU5F1* enhancer and generated sets of 63 sequences with an increasing number of prompt sequences. We calculated the AMS for each of the 879 JASPAR non-redundant vertebrate motifs across each generated set and observed that when there is only 1 sequence in the prompt, the generated sets do not have a strong POU5F1::SOX2 AMS (Figure 4D, blue line). However, once there are 3 or more homologs in the prompt, the generated sets have POU5F1::SOX2 AMS scores greater than 5 (Figure 4D, blue line). Altogether, these results suggest that larger prompt sizes lead to more successful enhancer designs, however, the success rate saturates when more than 63 sequences are used.

### Sorting homologs by length during training enables EnhancAR to generate shorter enhancer sequences while preserving predicted function

The ability to design short enhancers is important for applications ranging from gene therapy to synthetic biology, due to size restrictions in viral vectors (Wu et al., 2010) and minimizing unintended regulatory information (Artemyev et al., 2024). One such strategy used to design short biological sequences involves taking existing sequences with a target function and removing letters to identify the minimal substring that retains the function of the original sequence (Baron et al., 2025). We present an alternative strategy for shortening enhancers, which biases EnhancAR to generate shorter sequences that preserve the function of a prompt, which we define as enhancer shortening. We reasoned that training EnhancAR on length sorted homolog families would allow it to design shorter enhancer sequences because the model would learn the additional constraint that the next sequence in a set must be shorter than the preceding sequence. We trained an EnhancAR-sorted model on the same training set, where the only difference was sorting the homologs by decreasing length. To evaluate whether EnhancAR-sorted can shorten enhancers, we prompted the model with the same 500 HepG2, K562, and WTC11 specific enhancer homolog families we used to assess cell type specific enhancer design in the previous section, sorted by descending length, and generated 10 enhancers for each prompt. Using the generated sequence with the highest predicted activity in the target cell type, we compared the length of the generated sequence with the shortest prompt sequence and observed that 85% (1,260/1,476) of the generated sequences were shorter than the shortest prompt sequence (Supplementary Figure 5A-C, right). In comparison, only 0.02% (3/1,476) of the EnhancAR generated sequences were shorter than the shortest prompt (Supplementary Figure 5A-C, left).

Having demonstrated that EnhancAR-sorted generates shorter sequences than its prompt, we next asked whether these shortened sequences maintained the activity of the target human enhancer. We focused on prompts where the human sequence had predicted activity of greater than 2 log_2_ fold change (log_2_ FC) in the target cell type. We designated a design as successful when it had predicted activity greater than 2 log_2_ FC while also decreasing in length compared to the human sequence. Of the shortened designs, 38% (166/436) of HepG2, 43% (185/434) of K562, and 42% (161/384) of WTC11 designs were successful (Figure 5A-C). This contrasts with the shortest homologs, where 20% (88/436) of HepG2, 18% (78/434) K562, and 21% (81/384) of WTC11 homologs were successful. Notably, some of the designs were stronger than their human enhancer, even though they were shorter (Figure 5A-C, filled yellow points above the horizontal red line). On average, successful designs were about 20% shorter than the target human sequence. The largest decrease in sequence length while maintaining predicted human activity was seen for a K562 enhancer, which shrunk from 200 bp to around 53 bp (Figure 6D). Motif scanning for the GATA1::TAL1 motif found a strong match in both the human and generated sequences, which may explain how the shortened sequence maintained predicted activity. Additionally, the EnhancAR-sorted designs were often stronger than the shortest homolog sequences, with 88% (383/436) of HepG2, 90% (389/434) of K562, and 89% (342/384) of WTC11 generations having higher predicted activity. These results demonstrate that EnhancAR-sorted is capable of generating shorter enhancers while retaining predicted activity in the target cell type.

**Figure 5.**
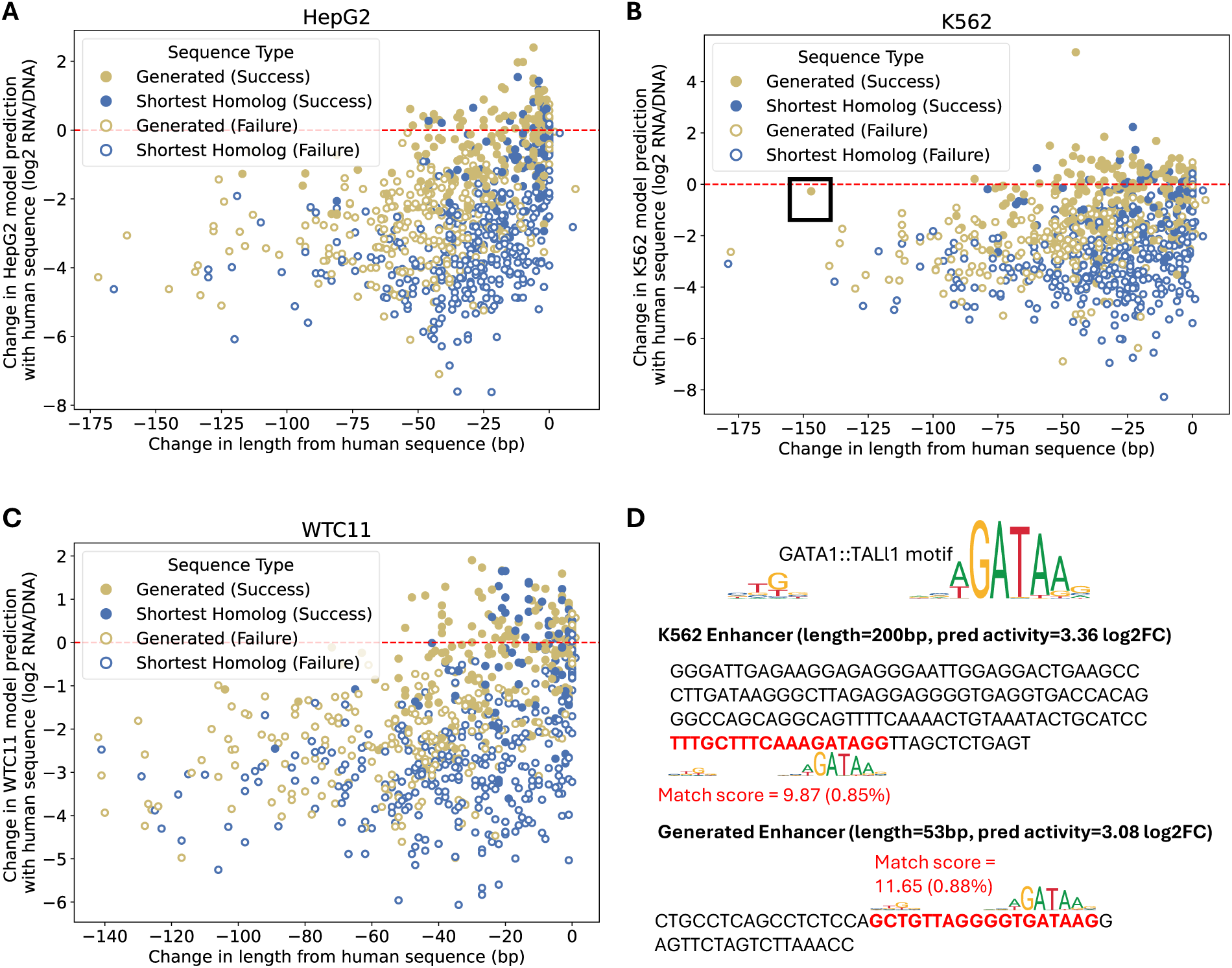
EnhancAR-sorted can shorten enhancers while preserving activity. (A-C) The change in length from the human sequence (bp) compared to change in HepG2 (A), K562 (B), and WTC11 (C) MPRALegnet model prediction with the human sequence for HepG2 (A), K562 (B), and WTC11 (C) prompted EnhancAR-sorted designs (yellow) and the shortest homolog sequence (blue). Design success (point fill) is defined as sequences with predicted activity >2 that are shorter than the human sequences (200 bp). The red dashed line represents no change in predicted enhancer activity between a sequence and the target human sequence. (D) The human enhancer and generated sequences for a predicted successful design. The GATA1::TAL1 motif from JASPAR (MA0140.1) is highlighted in red in both sequences.

## Discussion

We introduce EnhancAR, a generative model of enhancer evolution that leverages the relationship between homologs to generate novel enhancer sequences that share the function of their prompt sequences, even when function is not known. To our knowledge, this is the first time homology prompting with unaligned sequences has been used to design enhancers. EnhancAR was trained on a curated set of 1.7 million human enhancer homolog families spanning 1,888 cell types, which we believe is the largest dataset of homolog enhancer families to date. We demonstrate that when prompted by a homolog family for an enhancer, EnhancAR generates novel and diverse sequences that maintain sequence features from the prompt that are critical for regulatory function. Additionally, we show that EnhancAR can design cell type specific enhancers without further training, and that EnhancAR-sorted can shorten existing enhancers while maintaining predicted activity.

EnhancAR was designed based on the principle that homologous enhancers can differ in DNA sequence while preserving overall function, and that by comparing these shared features important for function can be identified. Thus, we trained EnhancAR on sets of unaligned homolog sequences, rather than treating each homolog as an individual. We reasoned that the long context of EnhancAR, enabled by the use of the Jamba architecture (Lieber et al., 2024), would permit EnhancAR to identify shared features across many homologs without using MSAs as input, enabling the model to learn relationships without bias from the alignment. Consistent with this hypothesis, we find that as we increased the number of homolog sequences in the prompt, enhancer designs showed on average higher predicted activity (Figure 4E, Supplementary Figure 4B/D). Similar observations have been observed in protein language models, where increasing MSA depth leads to more confident generations (Truong & Bepler, 2023).

As EnhancAR is designed to learn the regulatory logic of the prompted homolog family unsupervised, it could in theory design enhancers for any context, assuming it has an example sequence and its homologs. This contrasts with cell type prompted models, which require training on large datasets of thousands of examples to learn generalizable rules (DaSilva et al., 2026; Lal et al., 2024). Indeed, we demonstrated that EnhancAR can design sequences with specific activity in the target cell type by prompting EnhancAR with homologs for enhancers whose human sequence has the target specificity (Figure 4B). However, the more interesting application of generative enhancer models is not to design enhancers for lab defined cell lines, but rather to define enhancers for cells in contexts that are difficult to assay or do not have sufficient examples. For example, transient cell types during development are not stable and can be difficult to isolate, making them challenging to assay. Additionally, these cells often have overlapping regulatory logic with cells along the same lineage (Lettice et al., 2012; Oh et al., 2026), making it difficult to design with specificity. These represent real targets for applications such as gene therapy, which are suitable candidates for EnhancAR.

A limitation for the use of EnhancAR to design enhancers is that because it generates sequences that resemble real homologs for a homolog family, generations are expected to overlap with a similar distribution of function to the homologs, who may have lost enhancer function (Villar et al., 2015). Indeed, our results demonstrated that generations for HepG2, K562, and WTC11 enhancers displayed similar predicted activity and specificity to homologs, rather than the initial human sequences (Figure 4B) (although it remains unclear how much of this observation is explained by drift in homolog function, versus the MPRALegNet model being specific to human sequences). This is a trade off with our approach for the design of enhancers, as not all homologs will conserve the function of the human sequence. Thus, homologs act as an upper bound for EnhancAR’s enhancer designs. Future iterations may deal with this issue through the incorporation of phylogenetic relationships between homologs as done with other deep learning models (Karollus et al., 2024; Lim & Blanchette, 2020). EnhancAR could be extended to include species tokens (Brixi et al., 2025), aiding the model to learn about phylogeny. During inference, generations could be made to match a certain evolutionary distance by prompting with the species token, allowing EnhancAR to design a sequence for that species or evolutionary distance.

At inference time, EnhancAR consistently generated sequences of length 512 bp, regardless of the length of the sequences in the prompt (Supplementary Figure 2A). This observation is consistent with the maximum length constraint of 512 bp used during training and inference. We hypothesize that because the sequences in a prompt are not ordered by length, EnhancAR learns a weak probability of termination at each position. This is supported by our observation that when the maximum token generation size is set to 1024 bp, the probability of a separator token in the 1,000 held-out homolog families generations remains near zero until the 512 bp limit is reached and the probability quickly spikes (Supplementary Figure 2C). Additionally, EnhancAR-sorted, which is trained on increasingly shorter sequences, does not have this bias (Supplementary Figure 2D), suggesting that it is learning when to terminate generation. To mitigate this termination bias, EnhancAR generations were trimmed from the right-hand side to match the length of the longest prompt sequence. Future iterations of EnhancAR should aim to improve the capacity to learn when to terminate the generation.

While we have demonstrated that EnhancAR learns the importance of conserved motifs (Figure 3E/G), the presence of motifs is just one aspect of the cis-regulatory code; higher-order rules that govern interactions between transcription factors (e.g., motif spacing, orientation) are also important for enhancer function (Jindal & Farley, 2021). It is unclear whether EnhancAR is capturing these higher-order rules in the prompt sequences. In both genomic language models and natural language models, analysis of layers, such as the mamba, transformer, and mixture of expert layers (Gu & Dao, 2024; Vaswani et al., 2023), are used to interpret how these models are making predictions (Consens et al., 2023; Veiner & Supek, 2026). Similarly, the layers of EnhancAR could be investigated during next token prediction to understand the regulatory logic learned by EnhancAR. For example, if two motifs within a sequence are required for enhancer function, the internals of the model can be probed to determine whether higher attention is given between pairs of these motifs across homologs compared to background sequence. Subsequent experimental assays would validate whether the learned interactions are biologically relevant. This would shift EnhancAR from being a design tool to a method for understanding the cis-regulatory code.

## Methods

### Preparing data for EnhancAR training

To train EnhancAR, we downloaded candidate human proximal and distal enhancers from the ENCODE SCREEN V4 website (https://screen.wenglab.org/downloads), which includes 1,718,669 candidate enhancers (Moore et al., 2026). These regions are anchored on DNase Hypersensitivity Sites (DHS) and stratified based on a combination of measurements including DNase-seq, ATAC-seq, H3K4me3, H3K27ac, and CTCF across 1,888 human cell types, along with distance to nearest transcription start site (TSS).

Homologous sequences for each enhancer were identified in 240 non-human species from the Zoonomia 241-way vertebrate alignment (Zoonomia Consortium, 2020). Briefly, HAL (Hickey et al., 2013) and HALPER (Zhang et al., 2020) were used to identify homologs given human (hg38) enhancer coordinates. If multiple homologs were identified in another species, the first was arbitrarily selected. Altogether, 233,158,475 homologs were identified, with an average of around 135 homologs per enhancer. Each sequence and corresponding homologs were aligned using MAFFT (Katoh et al., 2002) with the adjust direction option. This was done to ensure homologous sequences were in the same orientation during training. Past this step, gap tokens representing the alignment are stripped from sequences.

To prepare unaligned homologous sets for training, we sampled up to 64 sequences per enhancer homolog family, prioritizing diversity by maximizing the Hamming distance. Then, we concatenated the sequences into a single string of characters, using slashes (/) to separate sequences and curly braces ({ }) to denote the start and end of the set. Finally, these sequences were tokenized using character level tokenization for input to the model.

The enhancer homolog sets were split into training, validation, and test sets. The test set contains all enhancers from chr3 and chr19. For the validation set, 1,000 of the training set MSAs were held out.

### EnhancAR architecture and training

All EnhancAR models are 170M-parameter models based on the Jamba hybrid transformer-state space model architecture (Lieber et al., 2024). We used the same architecture previously used for autoregressive protein sequence modeling by Dayhoff (Yang et al., 2025). Briefly, every 8th layer is a transformer module, with the remainder being Mamba modules (Gu & Dao, 2024). Every other Mamba module uses a mixture of experts (MoE) with 16 experts instead of a multilayer perception layer. All Mamba layers have state dimension 16, convolution dimension 4, and expansion dimension 2. In total, the architecture has 24 layers with model dimension 256, intermediate layer widths of 1024 inside each transformer block, 16 attention heads per transformer, and 8 key-value heads per transformer, resulting in 170,298,096 total parameters.

EnhancAR models are trained on unaligned homologous sets using an autoregressive cross-entropy loss with teacher forcing. We trained models on 8 MI300X GPUs for 50,100 steps, with a per-GPU batch size of 32. We used an inverse square root scheduler with a warm-up of 1,000 steps, max learning rate of 4e-4, and a MoE auxiliary loss weight of 0.1. Sequences were flipped at probability 0.5. For all models, we use the Adam optimizer (Kingma & Ba, 2017) with betas (0.9, 0.999). For EnhancAR-sorted only, we sorted sequences within homologous families from longest to shortest.

### Unconditional sequence generation

For the unconditional generation analysis, we unconditionally generated 1,000 sequence sets with 63 sequences per set using EnhancAR. Briefly, EnhancAR was prompted with only the homolog set start token ({), and was allowed to generate tokens A, T, C, G, and sequence separator (/). The generation was terminated either by the insertion of a sequence separator token or the maximum length of 512 bp was reached. If 512 bp was reached, a sequence separator token was inserted after the final nucleotide. The generated sequences were then fed back into EnhancAR as consecutive prompts, iteratively generating new nucleotides based on the previous generations.

### Conditional sequence generation using the test set

For the held-out enhancer homolog set analysis, we randomly sampled 1,000 enhancer homolog families from the test set as prompts to evaluate EnhancAR. We filtered out any homologs whose length was more than 2 standard deviations from the mean homolog length for a given homolog family to remove outliers. For each enhancer homolog family, we sampled up to 63 homologs and prompted EnhancAR with the homolog sequences separated by the sequence separator token, followed by a sequence separator token at the end. EnhancAR was allowed to generate tokens A, C, T, G, and the sequence separator token. The generation was terminated either by the insertion of a sequence separator token or the maximum length of 512 bp was reached. This process was repeated 63 times for each enhancer homolog family, using the same sampled homologs. As a heuristic, generations were trimmed from the right side so that they were no more than the maximum homolog length of the prompt.

### Measuring the novelty of sequences with training

To assess sequence similarity between the generated sequences and the training dataset, we performed a BLASTN (Camacho et al., 2009) search of all generated sequences against the training set (-evalue 1e-10). To establish a baseline for similarity in real homolog families, we performed BLASTN on the 1,000 sample enhancer homolog families from the held-out test set against the training set.

### Measuring the novelty of generations with the prompt

To assess sequence similarity between the generated sequences and their prompt, we performed glocal alignment with the BioPython (Cock et al., 2009) PairwiseAligner tool using BLASTN defaults (match_score=2, mismatch_score=-3, open_gap_score=-5, extend_gap_score=-2, target_end_gap_score=0) (Camacho et al., 2009) and reported the percent identity of the prompt sequence with the highest pairwise alignment score. We used glocal as generations could be different lengths from their prompt. The percent identity was calculated as the total number of matches (not including gaps) divided by the length of the smaller sequence. To establish a baseline for sequence similarity of a real homolog, we measured the sequence similarity of the held-out human sequence with their prompt. To establish a baseline for sequence similarity of unrelated sequences, we scramble the held-out human sequence and measure the sequence similarity with the prompt. Finally, as a comparison for a naïve generation approach, we compute the consensus sequence of each prompt and compare the sequence to their prompt.

### Measuring diversity of sequence sets

To investigate the sequence similarity of sequences within generated sequence sets, we performed glocal alignment with the BioPython (Cock et al., 2009) PairwiseAligner tool using BLASTN defaults (match_score=2, mismatch_score=-3, open_gap_score=-5, extend_gap_score=-2, target_end_gap_score=0) (Camacho et al., 2009), selected the highest scoring alignment for each pair, and reported the average percent identity of across all sequences in a set. The percent identity was calculated as the total number of matches (not including gaps) divided by the length of the smaller sequence. To establish a baseline of diversity for real enhancer homolog sets, we measured the diversity of the 1,000 sampled enhancer homolog families from the held-out test set. To establish a baseline for diversity in the absence of sequence homology, we randomly shuffled the nucleotides in each sequence in the 1,000 sampled enhancer homolog families from the held-out test set.

### Calculating average motif scores

We calculate the average motif scores based on a previous approach (Alam et al., 2023). We first downloaded the JASPAR 2024 non-redundant vertebrate motif set from JASPAR (Rauluseviciute et al., 2024). This dataset contains a reduced set of motifs, totaling 879 transcription factor motifs. Using the lightmotif python package (https://github.com/althonos/lightmotif), sequences in each set were scanned for the highest scoring match to each motif, which were then averaged to calculate the average motif score. This resulted in a vector of 879 average motif scores for each sequence set. To normalize the average motif score for a motif, the calculations were repeated for 100 scrambled versions of the motif. A Z-score was then calculated for the raw average motif scores.

### Comparing average motif scores

To investigate whether unconditionally generated sets shared motifs, we calculated the average motif scores for the unconditionally generated sets. For comparison, we also computed the average motif scores for the 1,000 sampled enhancer homolog families from the held-out test set. Additionally, to establish a baseline where there is no homology between sequences, we created new sequence sets by shuffling sequences across unconditionally generated sequence sets. Similarly, we created new sequence sets by shuffling sequences across the 1,000 sampled enhancer homolog families from the held-out test set.

### UMAP of average motif scores

To define the functional space of enhancers using motifs, we calculated average motif score representations for 50k randomly sampled training homolog sets (excluding *Mus musculus* and *Homo sapiens*) and then fit using UMAP (metric=cosine, n_neighbors=15, min_dist=0.1, random_state=42). Then, we projected the average motif score representations for the unconditionally generated sets and 1,000 held-out test set homolog sets into the UMAP space. For plotting, we estimated a 2D kernel density over the training-set UMAP coordinates and rendered it as a background density map. We overlaid scatter points for the held-out test set, and unconditional generations.

### UMAP of Enformer embeddings

To define the functional space of enhancers using predictive models, we calculated Enformer (Avsec et al., 2021) embeddings for 50k randomly sampled sequences (excluding *Mus musculus* and *Homo sapiens*) from the previous sampled 50k randomly sampled training homolog sets. We used sequence embeddings from the PyTorch (Paszke et al., 2019) implementation of Enformer as a fixed feature space to compare real and generated enhancer sequences. For each input sequence, we constructed an Enformer-length input of 196,608 bp by one-hot encoding nucleotides (A/C/G/T; all other characters mapped to N) and symmetrically zero-padding shorter sequences to center them within the 196 kb window. We then ran Enformer in inference mode and extracted the per-bin embedding track returned by the model, which has 896 bins spanning 114,688 bp at 128 bp stride. To obtain a single vector per sequence, we averaged the embedding vectors from the four central bins (center ±2 bins covering 512 bp), yielding a fixed-dimensional representation for each sequence.

We fit the Enformer embeddings for the 50k training homolog sequences using UMAP (n_neighbors=30, min_dist=0.1, random_state=42). Then, we projected the Enformer embeddings for randomly selected sequences from the unconditionally generated sets and 1,000 held-out test set homologs into the UMAP space. For plotting, we estimated a 2D kernel density (Gaussian KDE; Scott bandwidth) over the training-set UMAP coordinates and rendered it as a background density map (values below 1% of the maximum density masked). We overlaid scatter points for the held-out test set, and unconditional generations.

### Evaluating motif composition for specificity

To assess the specificity of generated sequences and their prompts, we compared the Pearson correlation of the average motif score representations between all the generated homolog sets and the prompts.

### Evaluating motif composition for importance

To measure whether the generated sequences preserved the important motifs of their prompts, we compared motifs with an average motif score greater than 5 in the generated sequences and the prompts. The precision and recall were calculated using the following definitions:

- False negatives: motifs whose AMS > 5 in the prompt but AMS < 5 in the generated sequences
- False positives: motifs whose AMS < 5 in the prompt but AMS > 5 in the generated sequences
- True negatives: motifs whose AMS < 5 in the prompt and AMS > 5 in the generated sequences
- True positives: motifs whose AMS > 5 in both the prompt and generated sequences

We also measured the average precision and recall of generations with all other prompts to assess specificity. For this analysis, we restricted prompts to only those with at least 2 motifs with Z-scores greater than 5.

To establish a baseline for a model that randomly mutates prompt sequences, we generated a control set by mutating residues at a mutation rate of 15% in each prompt sequence and measured the precision and recall with the prompt and all other prompts.

### Generating cell type specific enhancers

We downloaded cell type specific enhancers in HepG2, K562, and WTC11 cell lines from a recent publication that were tested with an MPRA (Agarwal et al., 2025). All these enhancers were 200bp long. Each enhancer had been experimentally measured in each cell type, and an element specificity score (ESS) was calculated to quantify the cell type specificity of each enhancer. We selected the 500 most specific enhancers by the highest ESS in each cell type with at least 64 sequences in the homolog family. Data was processed in the same manner as the training data, identifying 68,420 homologs for HepG2 peaks (mean 138 per family), 64,455 homologs for K562 peaks (mean 130 per family), and 63,171 homologs for WTC11 peaks (mean 127 per family). For each homolog family, we generated 10 sequences with EnhancAR. As a heuristic, we trimmed generations from the right side so that they are no more than the maximum homolog length of the prompt.

### Measuring motif occurrence

We used SEA (Bailey & Grant, 2021) from the MEME-suite to identify enriched motifs from the JASPAR 2024 Non-Redundant Vertebrate set (Rauluseviciute et al., 2024) in the generated and prompt sequences using default settings, focusing on cell type specific motifs for each cell line (Agarwal et al., 2025). Enrichment values of 1 are filled in for any value that does not meet the default Q-value threshold, as these are not reported be SEA.

### MPRALegNet Model

Joint MPRALegNet models (test1_val9) trained on HepG2, K562, and WTC11 MPRA datasets were downloaded from Zenodo (https://zenodo.org/records/10558183) (Agarwal et al., 2025). These models were trained to predict log2(RNA/DNA) activity measurements from 230 bp DNA sequences. Enhancer activity was predicted for each sequence in each of the cell types for both forward and reverse strands. The average predicted activity between forward and reverse strands is reported. Sequences shorter than 230 bp were embedded into a random 230 bp DNA sequence prior to prediction. Sequences longer than 230 bp were trimmed on the right side.

### Investigating the number of sequences in prompts

To evaluate the impact of increasing the number of sequences in the prompt, we selected the 273 HepG2, 262 K562, and 242 WTC11 enhancer homolog families from the previous 500 enhancer homolog families that had at least 152 homologs. We prompted EnhancAR with increasing numbers of sequences in each prompt, starting from no sequences (unconditional) to 151 sequences. We scanned each generation using lightmotif with P-value cutoff of 0.0001 for matches to the cell type specific factors POU5F1::SOX2 (JASPAR MA0142.1), HNF4A (HOCOMOCO12 HNF4A.H12CORE.0.PS.A) (Vorontsov et al., 2024), and GATA1::TAL1 (JASPAR MA0140.3). We calculated the proportion of generations at each number of prompt sequences with at least one match to each motif. As a comparison, we also measured the proportion of each motif in scrambled human sequence, human sequences, and homolog sequences. Additionally, we predicted the HepG2, K562, and WTC11 MPRA activity for each sequence using the MPRALegNet models previously mentioned.

### Investigating the distal POU5F1 enhancer

To investigate an example of a well-studied enhancer, we analyzed the core of the human distal *POU5F1* enhancer (hg38, chr6: 31173021-31173220). We extracted homologs for 227 mammalian species from the Zoonomia 241-way mammalian alignment. First, we generated 63 sequences using the same 63 sampled sequences from the *POU5F1* homolog family and calculated the POU5F1::SOX2 average motif score in the prompt and the generated set. Then, we generated 63 sequences with an increasing number of sequences in the prompt and calculated the average motif scores for all 879 JASPAR non-redundant vertebrate sequences for each generated set.

### Shortening enhancers using EnhancAR-sorted

We sorted the 500 K562, HepG2, and WTC11 enhancer homolog families by descending sequence length and prompted EnhancAR-sorted to generate 10 enhancers per prompt, holding out the human sequence. Using the MPRALegNet models, enhancer activity was predicted for each generation, all prompt sequences, and all held-out human sequences. We then filtered sequences where the human was predicted to have activity less than 2 log_2_ FC. For each enhancer, we used the best generation based on the highest predicted activity.

## Data availability

Data for training and evaluating EnhancAR and EnhancAR-sorted are available on Zenodo at https://zenodo.org/records/19052620.

## Code availability

Code for training EnhancAR and analyzing the data is available on GitHub at https://github.com/microsoft/enhancAR. The models and weights for EnhancAR and EnhancAR-sorted are available on Hugging Face at https://huggingface.co/aduncan94/EnhancAR and https://huggingface.co/aduncan94/EnhancAR-Sorted, respectively.

## Acknowledgements

We thank Cameron Dufault, Ami Sangster, Denise Le, and Rain Jin of the Moses lab for their helpful comments on the manuscript.

## Supplementary figures

**Supplementary Figure 1.**
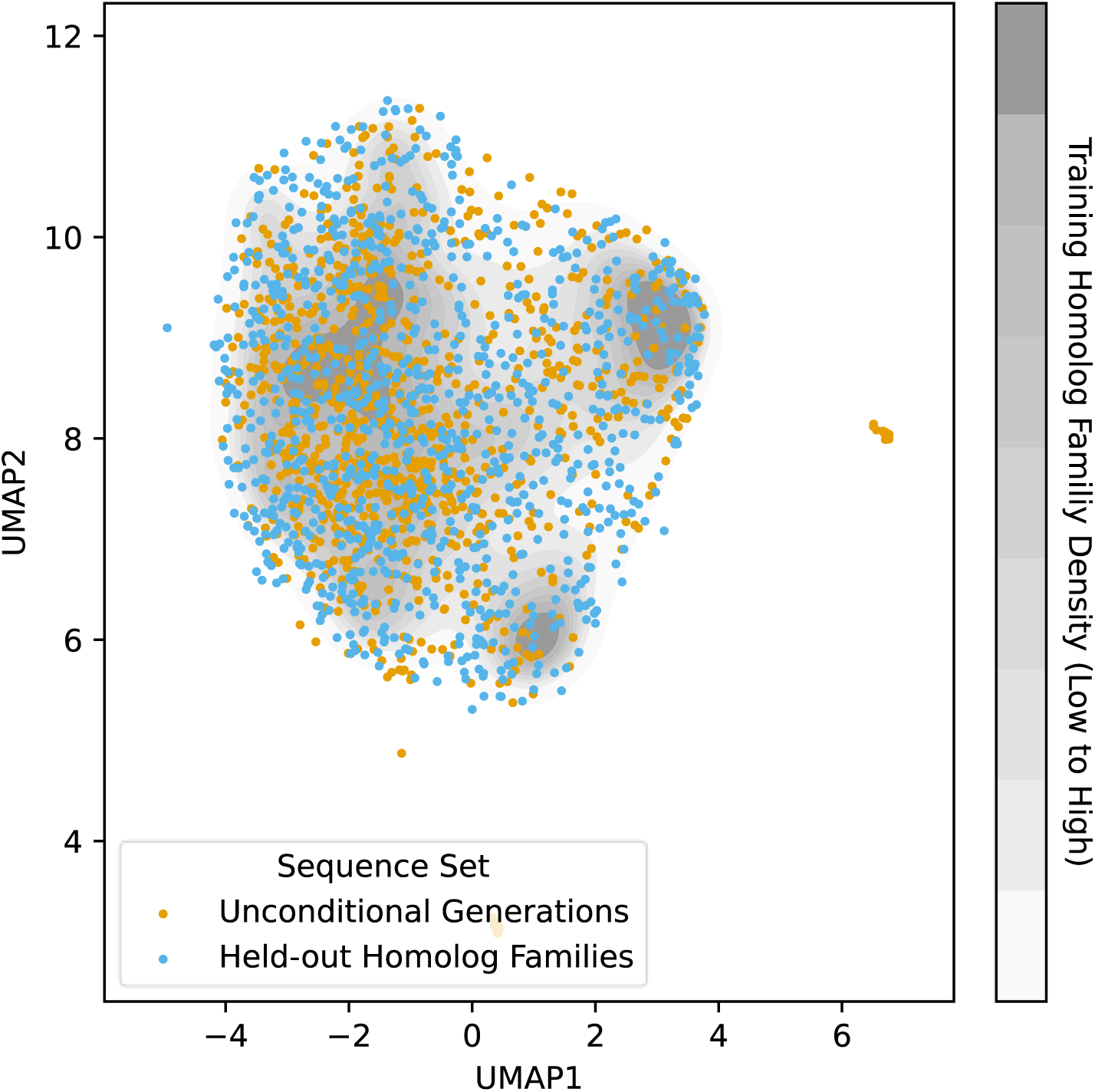
The average motif score representations of unconditionally generated sets stay near the mode of the training enhancer homolog family distribution. Scatterplot of the average motif score UMAP of training data (grey kernel density plot) (n=50,000), unconditional generations (yellow points) (n=1,000), and held-out homolog families (blue points) (n=1,000). The UMAP was fit to training dataset, and the unconditional generations and held-out homolog families were transformed using the UMAP model.

**Supplementary Figure 2.**
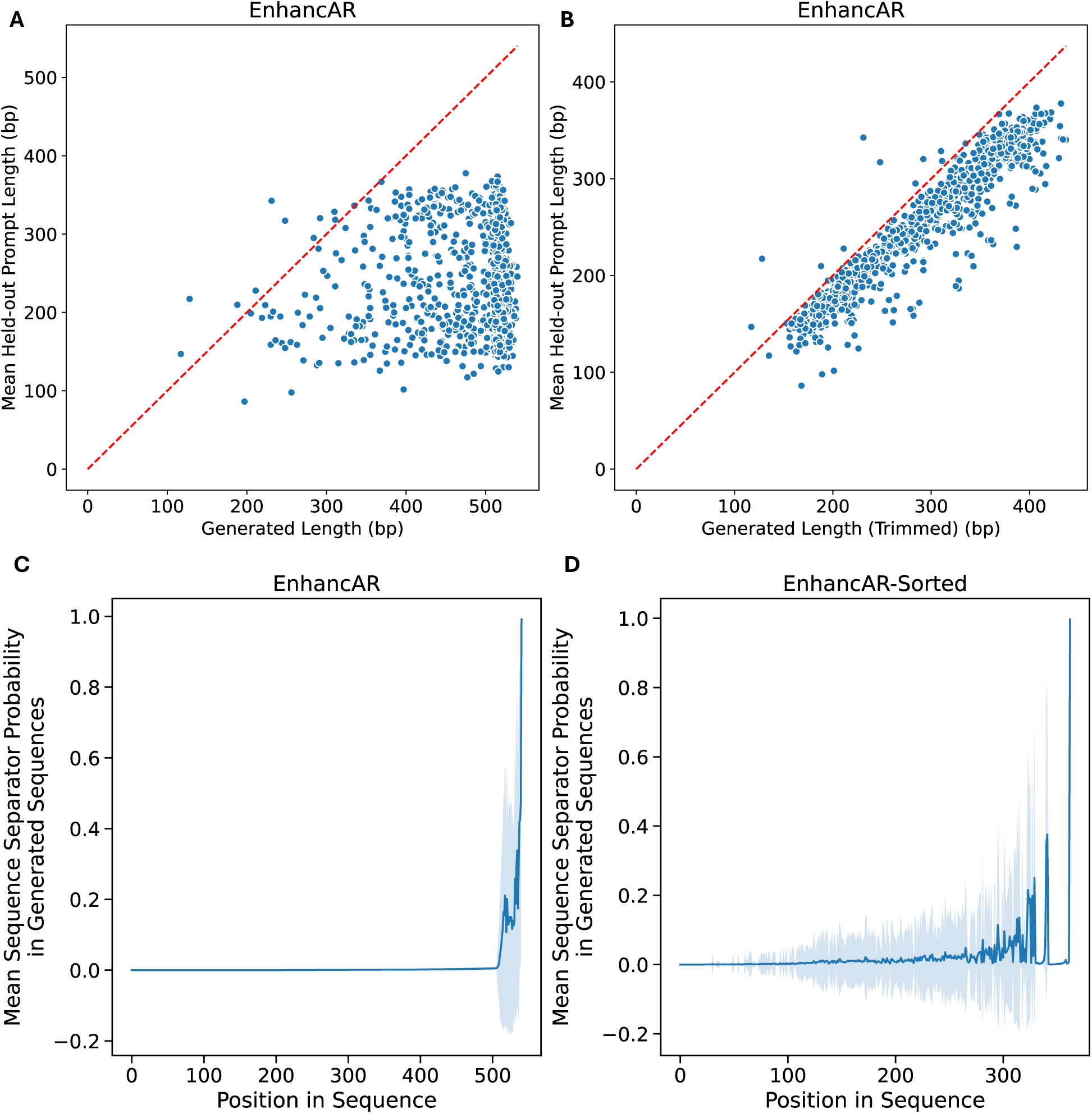
Test set prompted generations terminate near 512 bp training and inference limit. (A) The length of generated sequences (n=1,000) compared to their mean prompt length for EnhancAR held-out homolog family prompted generations. (B) The length of generated sequences (n=1,000), trimmed to their maximum prompt sequence length, compared to their mean prompt length for EnhancAR held-out homolog family prompted generations. (C) The mean sequence separator probability at each position across EnhancAR generations (n=1,000) for the held-out homolog families. (D) The mean sequence separator probability at each position across EnhancAR-sorted generations (n=1,000) for held-out homolog families. (C/D) The light blue fill represents the standard deviation of the sequence separator probability across the generations (n=1,000) at each position.

**Supplementary Figure 3.**
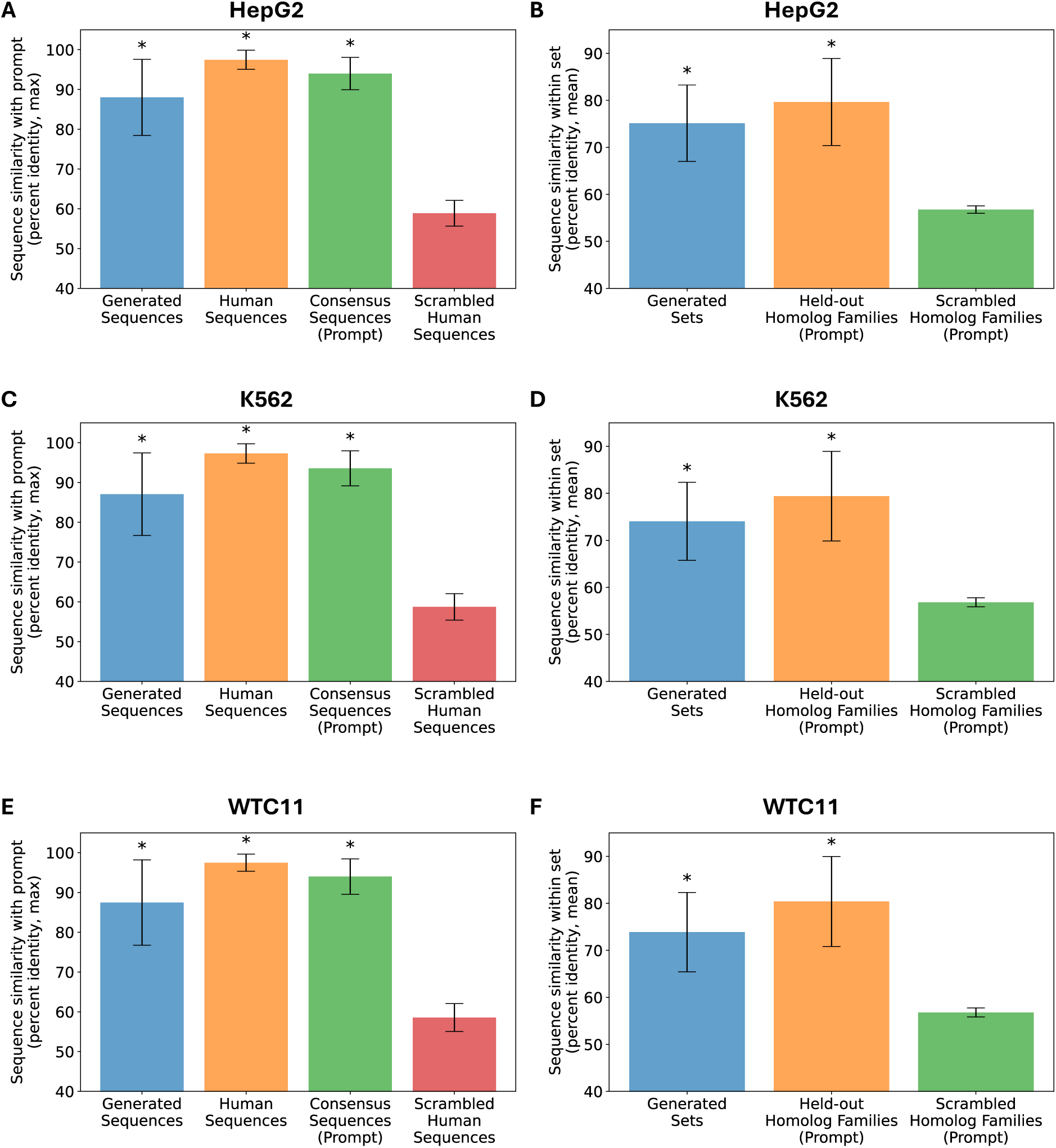
HepG2, K562, and WTC11 prompted generations are novel and diverse. (A/C/E) Bar plot showing the sequence similarity between sequences and their prompt, measured as the maximum percent identity, for EnhancAR generated sequences, human sequences, and scrambled human sequences for HepG2 specific enhancers (n=500) (A), K562 specific enhancers (n=500), and WTC11 specific enhancers (n=500). Significance was assessed by comparing groups to the scrambled human control using the Mann-Whitney U test. Error bars represent the standard deviation of the maximum percent identity. (B/D/F) Bar plot sequence similarity between sequences within a set, measured as the mean percent identity of the EnhancAR generated sequences, the prompt sequences, and scrambled prompt sequences for HepG2 specific enhancers (n=500), K562 specific enhancers (n=500), and WTC11 specific enhancers (n=500). Significance was assessed by comparing groups to the scrambled homolog sets using the Mann-Whitney U test. Asterisks represent significance (p-value < 0.05) after Benjamini-Hochberg correction. Error bars represent the standard deviation of the mean percent identity.

**Supplementary Figure 4.**
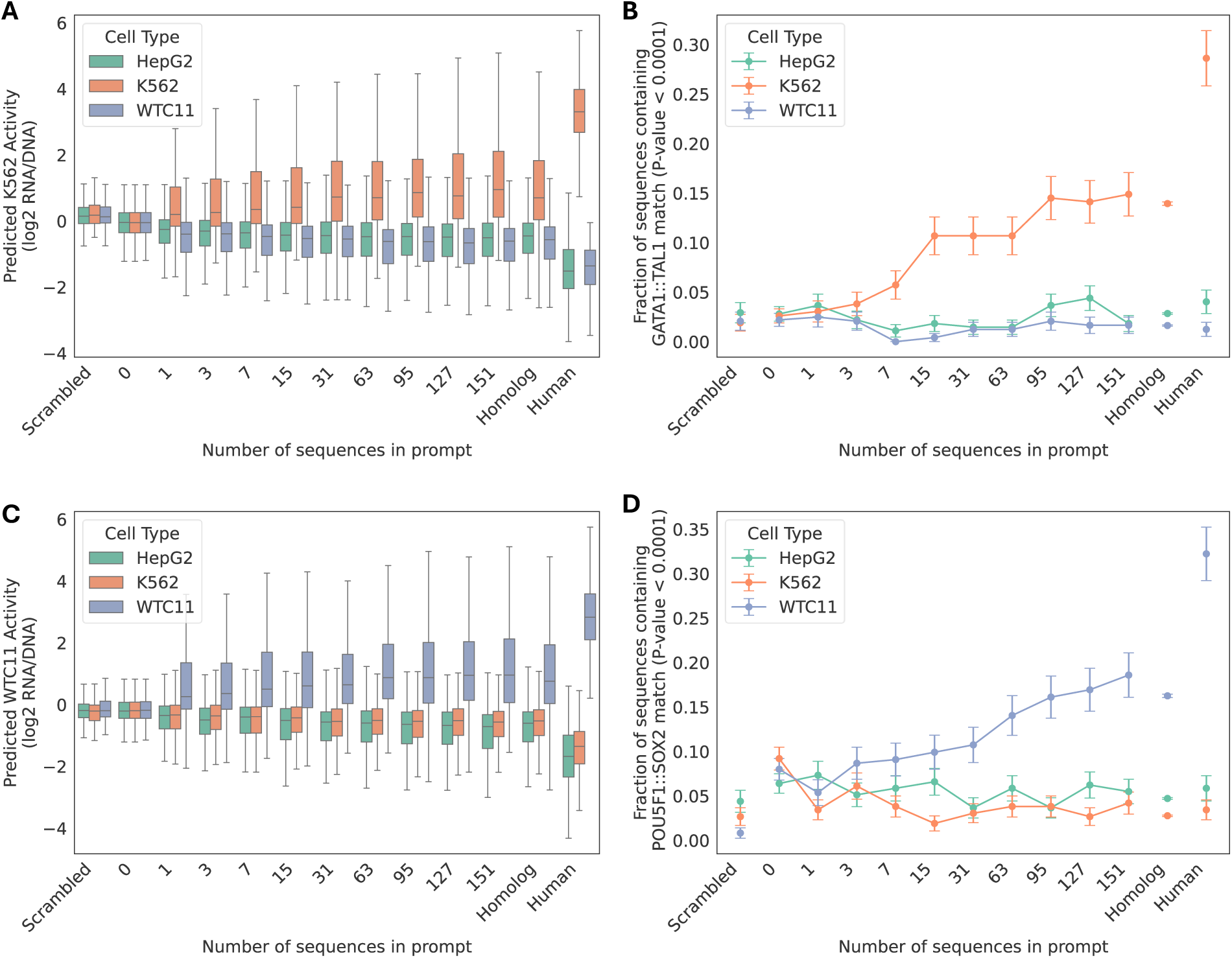
EnhancAR design success increases with prompt number. (A/C) Predicted K562 (A) and WTC11 (C) enhancer activity (log2(RNA/DNA)) using the MPRALegNet models is shown on the y-axis for the HepG2 (n=273), K562 (n=262), and WTC11 (n=242) prompted EnhancAR designs generated with an increasing number of sequences in the prompt. (B/D) The proportion of HepG2, K562, and WTC11 prompted EnhancAR designs generated with an increasing number of sequences in the prompt that contain a significant match to the K562 specific factor GATA1::TAL1 motif (B), and the WTC11 specific factor POU5F1::SOX2 JASPAR motif (D) using the lightmotif package (P-value <1e-4).

**Supplementary Figure 5.**
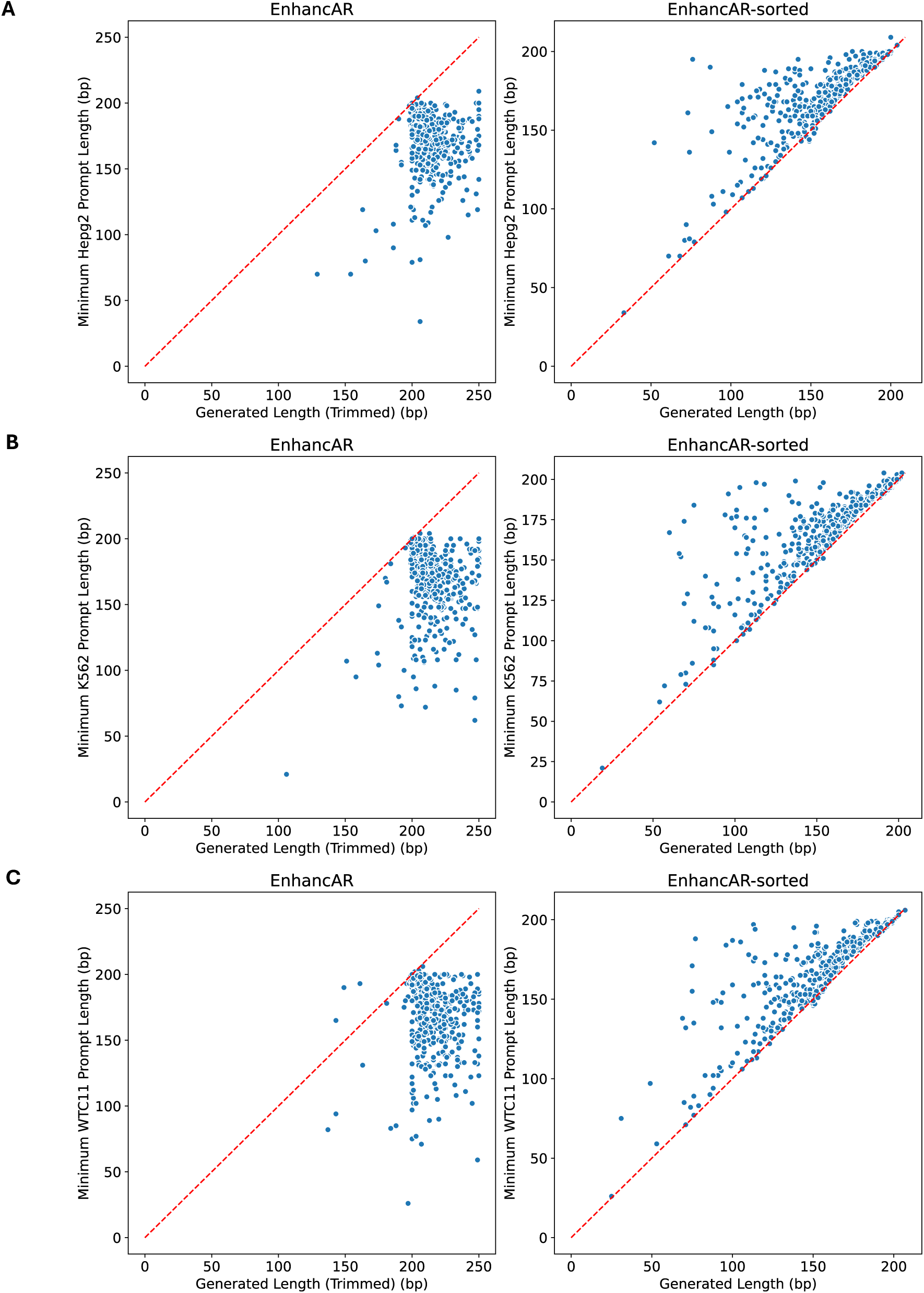
EnhancAR-sorted generations are often shorter than the sequences in their prompt. (A) The length of generated sequences (n=500) compared to the minimum prompt length for EnhancAR (left) and EnhancAR-sorted (right) for HepG2 prompted generations. (B) The length of generated sequences (n=500) compared to the minimum prompt length for EnhancAR (left) and EnhancAR-sorted (right) for K562 prompted generations. (C) The length of generated sequences (n=500) compared to the minimum prompt length for EnhancAR (left) and EnhancAR-sorted (right) for WTC11 prompted generations.

